# Vaginal squamous cell carcinoma develops in mice with *Arid1a* loss and gain of oncogenic *Kras*

**DOI:** 10.1101/2020.12.15.422959

**Authors:** Xiyin Wang, Mariana S. L. Praça, Jillian R. H. Wendel, Robert E. Emerson, Francesco J. DeMayo, John P. Lydon, Shannon M. Hawkins

**Affiliations:** Department of Obstetrics and Gynecology, Indiana University School of Medicine, Indianapolis, IN, USA; Deparment of Obstetrics and Gynecology, Federal University of Minas Gerais, Belo Horizonte, Minas Gerais, Brazil; Group of Health, Hospital Mater Dei, Belo Horizonte, Minas Gerais, Brazil; Department of Pathology and Laboratory Medicine, Indiana University School of Medicine, Indianapolis, IN, USA; National Institute of Environmental Health Sciences, Research Triangle Park, NC, USA; Department of Molecular and Cellular Biology, Baylor College of Medicine, Houston, TX, USA

**Keywords:** ARID1A, KRAS, vaginal squamous cell carcinoma, mouse model

## Abstract

Recent sequencing studies showed that loss-of-function mutations in *ARID1A* (AT-rich interactive domain 1a) were enriched in gynecologic malignancies. However, multiple mouse models with deletion of *Arid1a* did not exhibit gynecologic malignancy. Oncogenic *KRAS* mutations are a common finding in endometrial cancers. However, expression of oncogenic Kras (*Kras^G12D^*) in the uterus was not sufficient to develop endometrial cancer. These results suggest that both ARID1A deletion and oncogenic KRAS require additional hits before driving gynecologic malignancy. To determine the role of the combination effects of deletion of *Arid1a* and oncogenic *Kras, Arid1a^flox/flox^* mice were crossed to *Kras^Lox-Stop-Lox-G12D/+^* mice using progesterone receptor Cre (*Pgr^Cre/+^*). Survival studies, histology, and immunohistochemistry were used to characterize the phenotype. Hormone dependence was evaluated by ovarian hormone depletion and estradiol replacement. *Arid1a^flox/flox^*; *Kras^Lox-Stop-Lox-G12D/+^*; *Pgr^Cre/+^* (AKP) mice exhibited early euthanasia due to large vaginal tumors, which were invasive squamous cell carcinoma. Younger mice exhibited precancerous intraepithelial lesions that progressed to invasive squamous cell carcinoma with age. Immunohistochemistry supported the pathological diagnosis with abnormal expression and localization of cytokeratin 5, tumor protein P63, cyclin dependent kinase inhibitor 2A (CDKN2A or p16), and marker of proliferation Ki-67. Vaginal lesions in AKP mice were hormone dependent. Ovarian hormone deletion in AKP mice resulted in atrophic vaginal epithelium without evidence of vaginal tumors. Estradiol replacement in ovarian hormone depleted AKP mice resulted in lesions that resembled the squamous cell carcinoma in intact mice. AKP mice did not develop endometrial cancer. *Arid1a* deletion with *Kras^G12D^* expression drives invasive vaginal squamous cell carcinoma. This mouse can be used to study the transition from benign precursor lesions into invasive vaginal squamous cell carcinoma offering insights into progression.

## INTRODUCTION

Recent sequencing studies, including both TCGA (The Cancer Genome Atlas) PanCancer and MSK-IMPACT (Memorial Sloan-Kettering Integrated Mutation Profiling and Actionable Cancer Targets) have highlighted potentially impactful mutations across gynecologic cancers [1,2]. Worldwide in 2018, more than 1.3 million women were diagnosed with a gynecologic cancer, and over 528,000 women with gynecologic cancer died [3]. In silico analyses of multiple, large, publicly-available datasets have shown that a small subset of genes are mutated across uterine cervix, uterine corpus, ovary, vulva, or vagina cancers [4]. However, the functional role of each of these genes in female reproductive tract malignancy remains largely unknown. Further, models of these diseases would improve the understanding of disease pathogenesis, accelerate discovery of novel therapeutics, and improve lives of many women worldwide.

*ARID1A* (AT-rich interactive domain 1A) was one of eight genes whose mutation frequency was significantly higher in gynecologic malignancies over other cancers [4]. Loss-of-function mutations in *ARID1A* are frequent in endometriosis-associated ovarian cancers, endometrial cancers, and cervical cancers [1,2,4–11]. More than 40% of endometrial cancers have a mutation in *ARID1A* [7–9]. Out of human papilloma virus-negative cervical squamous cell carcinomas, 33% contained mutations in *ARID1A* [11]. *ARID1A* encodes a protein in the switch/sucrose non-fermentable chromatin remodeling complex, playing a role in transcriptional regulation and reprogramming [9,12,13]. ARID1A plays an important role in female reproduction and is ubiquitously expressed in the female reproductive tract [14,15]. Conditional deletion of *Arid1a* with the anti-Müllerian hormone receptor *2-Cre* resulted in subfertility due to abnormal placentation [14]. Conditional deletion of *Arid1a* with the progesterone receptor-Cre *(PgrC^re/+^)* or the lactotransferrin-Cre resulted in infertility due to endometrial dysfunction [15,16]. Deletion of *Arid1a* alone in the mouse female reproductive tract was not sufficient to drive cancer. Additional gene deletions were required for development of gynecologic malignancies [12,13,17].

Mutations in *KRAS* (KRAS proto-oncogene, GTPase) are frequent in gynecologic malignancies, being mutated in more than 10% of samples across multiple large datasets [4]. Oncogenic *KRAS* has been detected in up to 30% of endometrial cancers [18–20]. Increased expression of *KRAS* was also associated with low overall survival across gynecologic cancers [4]. Conditional expression of *Kras^G12D^* alone was insufficient to develop cancer in mice [21–24]. However, *Pten* (phosphatase and tensin homolog) conditional deletion in *Kras^G12D^* expressing mice was sufficient for gynecologic malignancy [22–26]. Similar to deletion of *Arid1a* alone, gain of oncogenic *Kras^G12D^* alone in the female reproductive tract was not sufficient to drive cancer and additional gene deletions were required.

Concurrent mutations in both *ARID1A* and *KRAS* are frequent in mesonephric-like Müllerian adenocarcinomas of the female reproductive tract [27]. Similarly, concurrent mutations are common in endometrial and cervical cancer, but rarer, yet present, in ovarian cancer [7,11,28,29]. We hypothesized that conditional *Arid1a* deletion in *Kras^G12D^* expressing mice using *Pgr^Cre^* would generate female reproductive tract cancer.

## MATERIALS and METHODS

### Animal husbandry and genotyping

Animal experiments were approved by the Indiana University School of Medicine Institutional Animal Care and Use Committee following the National Institutes of Health Guide for the Care and Use of Laboratory Animals. All mice were bred and kept under standard conditions. *Arid1a^flox/flox^; PgrC^re/+^* [15] and *Kras^Lox-Stop-Lox-G12D/+^* mice [23] or *Pgr^Cre/+^* [30] and *Kras^Lox-Stop-Lox-G12D/+^* mice [23] were crossbred and maintained on a C57BL/6J; 129S5/Brd mixed hybrid background to generate *Arid1a^flox/flox^; Kras^Lox-Stop-Lox-G12D/+^;Pgf^Cre^/^+^* (AKP), *Kras^Lox^-^Sto^p-^Lox^-^G12D/+^;PgrC^re/+^* (KP), and *Pgr^+/+^* (control) mice (Supplementary Figure S1). Mice were genotyped at 12-14 days of postnatal life from tail biopsies by PCR as described [14,31,32]. *Cre*-mediated recombination in *Arid1a^flox/flox^* mice removes exon 8, leading to loss of protein [33]. *Cre*-mediated recombination of *Kras^Lox-Stop-Lox-G12D/+^* mice removes the stop codon, resulting in expression of oncogenic *Kras^G12D^* [34]. *Pgr^Cre/+^* drives *Cre*-mediated recombination where progesterone receptor is expressed [30]. For survival studies, mice were caged, examined twice weekly, and euthanized at humane endpoints [32]. Postmortem tail clips were used to confirm genotyping [14,31,32], and mice with inconsistent genotypes were reassigned or removed.

### Histological analyses

Tissues were fixed, stained, and quantified as described [32]. The Lower Anogenital Squamous Terminology (LAST) project nomenclature was used to describe histology [35]. All histology was interpreted by a surgical pathologist (R.E.E.). Antibodies for immunohistochemistry are listed in Supplementary Table S1. Immunohistochemistry was performed as described [32].

### Serum analysis

Mice were anesthetized by isoflurane (Abbott Laboratories, North Chicago, IL), and blood was collected in microtainer tubes (Becton Dickinson, Franklin Lakes, NJ) by closed cardiac puncture. Serum was separated by centrifugation and stored at −20°C until use. The University of Virginia Ligand Assay and Analysis Core performed measurements of follicle stimulating hormone (FSH) and luteinizing hormone (LH) by radioimmunoassay.

### Steroid hormone treatment

Mice underwent ovariectomy (ov’ex) at six weeks. Mice were randomly divided into treatment groups: estradiol pellet (Innovative Research of America, Sarasota, Florida, 0.25 mg 17β-estradiol/60-day release pellet) or no pellet (sham), and tissues collected after 60 days.

### Statistical analysis

Log-rank (Mantel-Cox) test, Fisher’s exact test, Student t-test, and multiple t-test was performed using the InStat package for GraphPad Prism 8 (GraphPad, San Diego, CA). *P* < 0.05 was considered statistically significant.

## RESULTS

### Large vaginal tumors

Based on criteria for humane endpoints [32], AKP mice showed decreased survival compared to *Pgr^+/+^* (Figure 1A and Supplementary Figure S2). The median survival of AKP mice was 106 days. AKP mice (10 of 10 mice) exhibited gross, solid lesions protruding from the vagina, resulting in early euthanasia. These lesions were pink-red, with oblong to round growth on the outer part of the vagina (Figure 1B). Upon dissection, the gross lesions appeared to be confined to the vagina, with grossly normal ovaries, oviducts, and uterus (Figure 1C).

**Figure 1.**
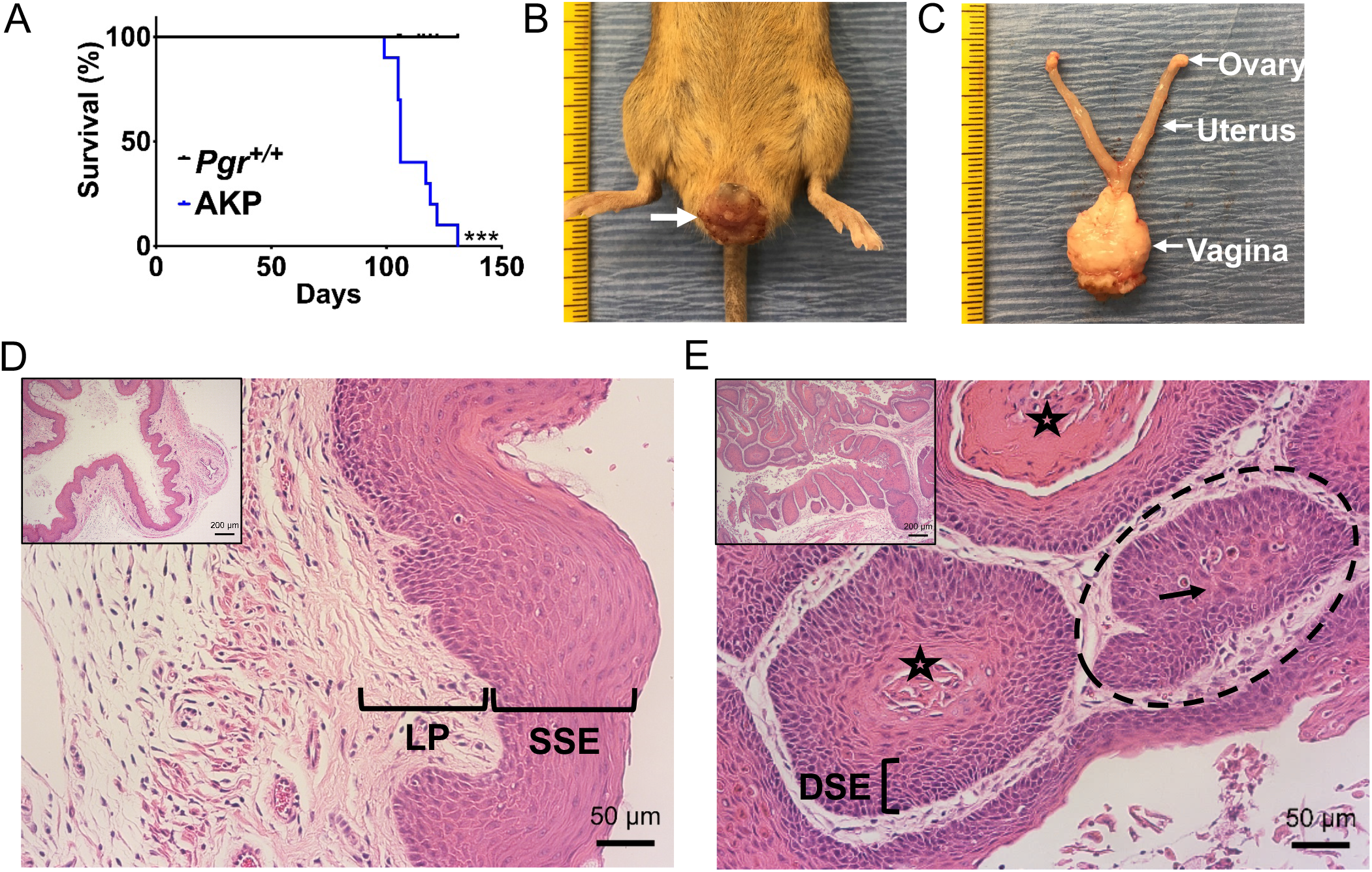
Poor survival and squamous cell carcinoma in AKA female mice. (A) Kaplan-Meier survival curves were analyzed by log-rank (Mantel-Cox) test. AKP mice have decreased survival compared to control mice. *** *P* < 0.001. *n* = 10 per group. (B) Gross vaginal tumor burden (white arrow) in AKP mice. (C) Gross lesions were localized to the vagina with grossly normal ovaries, oviducts, uterus, and cervix. Ruler, tick mark indicates 1 mm. (D) Control vagina showed normal keratinized stratified squamous epithelium (SSE) and lamina propria (LP). (E) AKP vagina exhibited squamous cell carcinoma lined by highly dysplastic squamous epithelium (DSE) with central keratinization (star) and nests of cells (dashed circle) with abundant eosinophilic cytoplasm (black arrow). Scale bars, low-power (insets), 200 μm; high-power, 50 μm. Hematoxylin and eosin staining.

*Pgr^+/+^* female mice had normal appearing vaginal histology with well-defined epithelial layers, including a keratinized outer layer of cells without nuclei, stratified squamous epithelium, and lamina propria (Figure 1D). AKP mice showed squamous cell carcinoma with central keratinization surrounded by highly dysplastic squamous epithelium as evidenced by the large, dark nuclei and disorganized cellular pattern, along with nests of cells with abundant eosinophilic cytoplasm (Figure 1E). Additional examples of squamous cell carcinoma are in Supplementary Figure S3, and individual mouse female reproductive tract histology is listed in Supplementary Table S2.

### Early Precancerous Lesions to Cancer

In women, invasive squamous cell carcinoma develops from precancerous lesions, including low grade squamous intraepithelial lesion (LSIL, also known as vaginal intraepithelial neoplasia 1, VaIN1) or high grade squamous intraepithelial lesion (HSIL, VaIN2 and VaIN3) [36]. As AKP mice aged, the penetrance of squamous cell carcinoma increased (Figure 2A). Squamous intraepithelial lesions were not found in *Pgr^+/+^* mice (Figure 2B). As early as eight weeks, 10% (1/10) of AKP mice exhibited LSIL. The nuclei in the epithelium were enlarged with variable size, irregular nuclear contours, and increased nuclear-to-cytoplasmic ratios (Figure 2C). Most frequently (7/10), AKP mice at eight weeks had HSIL. The epithelium exhibited a full-thickness proliferation of abnormal parabasal-like cells with loss of maturation and increased nuclear-to-cytoplasmic ratios (Figure 2D). As early as eight weeks, AKP mice (2/10) had invasive squamous cell carcinoma (Figure 2E). Additional magnification views are in Supplementary Figure S4. By 12 weeks, nearly 70% (16/23) of AKP mice developed squamous cell carcinoma (Figure 2A and Supplementary Table S3). At 16 weeks, 100% (11/11) AKP mice exhibited squamous cell carcinoma (Figure 2A and Supplementary Table S3).

**Figure 2.**
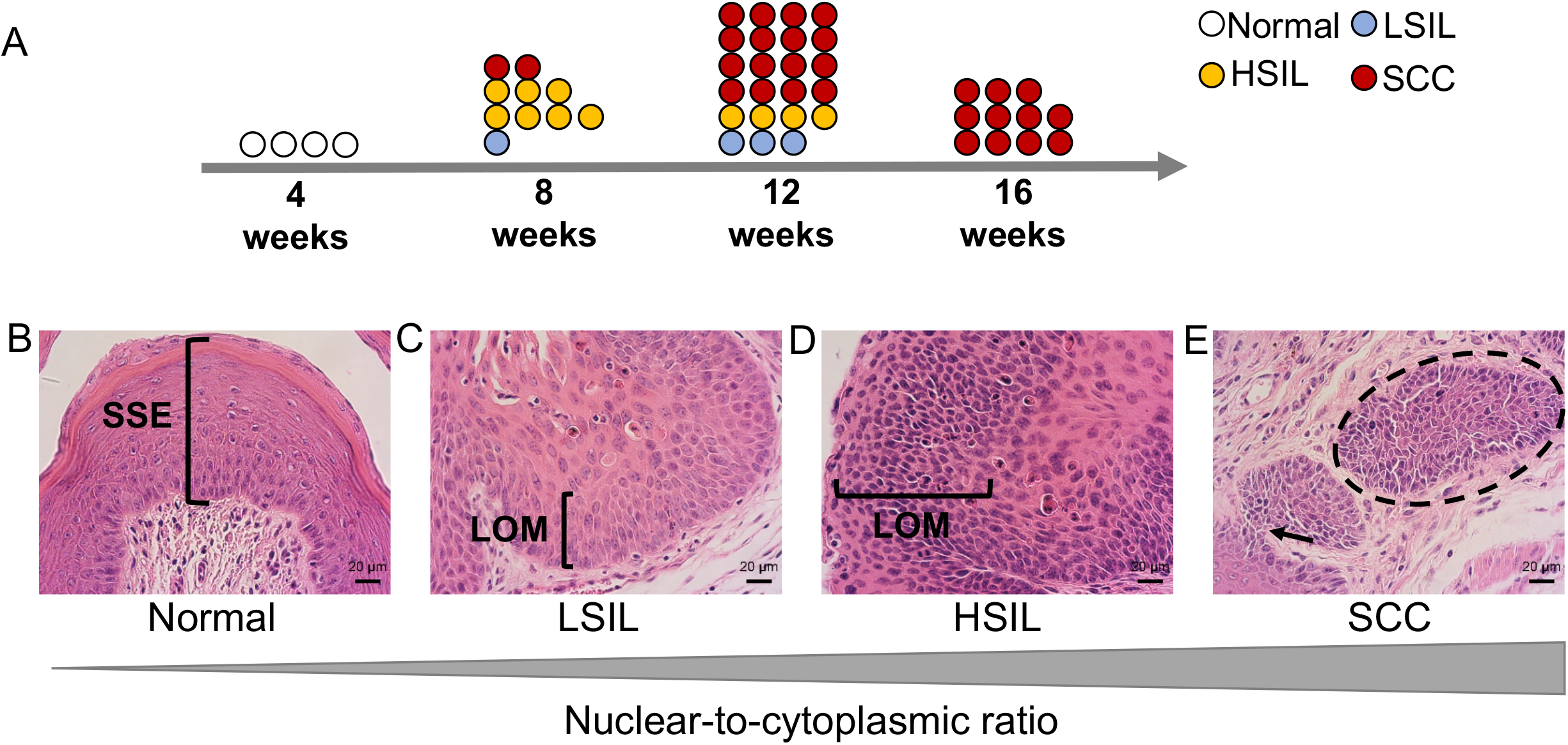
Squamous intraepithelial lesions and squamous cell carcinoma in AKP mice. (A) Schematic timeline of the progression of squamous intraepithelial lesions to squamous cell carcinoma in AKP mice. At 8 weeks, (B) *Pgr^+/+^* vagina showed normal stratified squamous epithelium (SSE). (C) Ten percent of AKP vaginas contained LSIL with increased nuclear-to-cytoplasmic ratios and loss of maturation (LOM) in the lower epithelium. (D) Seventy percent of AKP vaginas exhibited HSIL with cell crowding, high nuclear-to cytoplasmic ratio, and LOM in at least two-thirds of the epithelium. (E) Twenty percent of AKP vaginas displayed squamous cell carcinoma with nests of cells (dashed circle) and highly dysplastic squamous epithelium invading through the stroma (black arrow). Scale bars, 20 μm. Hematoxylin and eosin staining.

Because previous studies examined the two-month time point for *Kras^lox-stop-lox-G12D/+^; Pgr^Cre/+^* (KP) mice [24], we examined KP mice for precancerous lesion development. At eight weeks, only one out of six KP mice exhibited HSIL (Supplementary Figure S5 and Supplementary Table S4). The frequency of abnormal squamous histology was higher in AKP mice than KP mice at eight weeks (Fisher’s exact test = 0.0014, *P* < 0.01).

### Atypical Lesions Outside the Vagina

*Pgr^Cre/+^* results in *Cre*-mediated recombination in the uterus, oviduct, and ovaries [30]. However, no malignancy was identified in the AKP mice besides the vagina. AKP mice had normal ovaries, with normal follicular development (Supplementary Figure S6). Consistent with normal follicular steroid hormone feedback, we observed no significant differences between control and AKP mice in follicle stimulating hormone (FSH) levels *(Pgr^+/+^:* 3.607 ± 0.363 ng/ml, n = 10; AKP: 4.952 ± 1.857 ng/ml, n = 10; unpaired, two-tailed Student t-test, *P* = 0.487) or luteinizing hormone (LH) levels *(Pgr^+/+^:* 0.230 ± 0.060 ng/ml, n = 12; AKP: 0.200 ± 0.031 ng/ml, n = 10; unpaired, two-tailed Student t-test, *P* = 0.666).

Infrequently (5/14 mice), the oviductal epithelium in AKP mice at 16 weeks contained atypical epithelium with nuclear stratification and multiple layers of epithelial cells with nuclear atypia in the form of nuclear enlargement and rounding (Supplementary Figure S6 and Supplementary Table S3). There was no difference in body weight between AKP and *Pgr^+/+^* mice at any time points (data not shown). Uterine weight per body weight in AKP mice at 16 weeks was heavier than control (AKP: 4.2+/0.29 mg/g; control: 3.2+/-0.29 mg/g; multiple t-test, *P* <0.05), but there was no significant difference in uterine weight at 12 weeks (AKP: 3.5+/-0.25 mg/g; *Pgr^+/+^:* 3.4+/0.35 mg/g; multiple t-test, *P* =0.88) or 8 weeks (AKP: 2.9+/-0.16 mg/g; *Pgr^+/+^:* 3.5+/-0.69 mg/g; multiple t-test, *P* =0.28). Endometrial hyperplasia was observed in 21% (3/11) AKP mice at 16 weeks with enlarged glands separated by minimal amounts of stroma (Supplementary Figure S7 and Supplementary Table S3). Nuclear atypia was also observed with loss of polarity and rounding of the nuclei (Supplementary Figure S7). Endometrial adenocarcinoma was not observed in AKP mice. There was no squamous cell carcinoma detected in the cervix of AKP mice (Supplementary Table S3).

### Molecular Characterization

Cytokeratin 5 (KRT5) is a useful stain for squamous differentiation [36,37]. The basal epithelial cells of *Pgr^+/+^* vagina showed intense perinuclear and moderate cytoplasmic KRT5 expression with little to no staining in the lamina propria (Figure 3A and Supplementary Figure S8). Intense cytoplasmic and perinuclear KRT5 staining was observed throughout the full thickness of the vaginal squamous cell carcinoma nests but sparing the underlying lamina propria (Figure 3A and Supplementary Figure S8). Intense nuclear staining for the basal epithelial cell marker tumor protein P63 (p63) [38] was demonstrated in the basal layer of the *Pgr^+/+^* vagina. Moderately intense nuclear p63 staining was observed across the full thickness of the nests of squamous cell carcinoma in AKP vaginas (Figure 3B and Supplementary Figure S8). Vaginal squamous cell carcinoma in women has been shown to have diffusely positive cyclin dependent kinase inhibitor 2A (CDKN2A also known as p16) staining [36,39]. The *Pgr^+/+^* vagina showed low intensity, low frequency nuclear staining for p16 that was limited to the basal layer. Nest of vaginal squamous cell carcinoma in AKP mice showed intense, diffusely positive p16 staining (Figure 3C and Supplementary Figure S8). Marker of proliferation ki67 (Ki67) staining in *Pgr^+/+^* mice was contained to a mostly single layer of basal epithelium of the vagina. Ki67 staining in AKP mice was shown in multiple layers in nests of squamous cell carcinoma (Figure 3D and Supplementary Figure S8).

**Figure 3.**
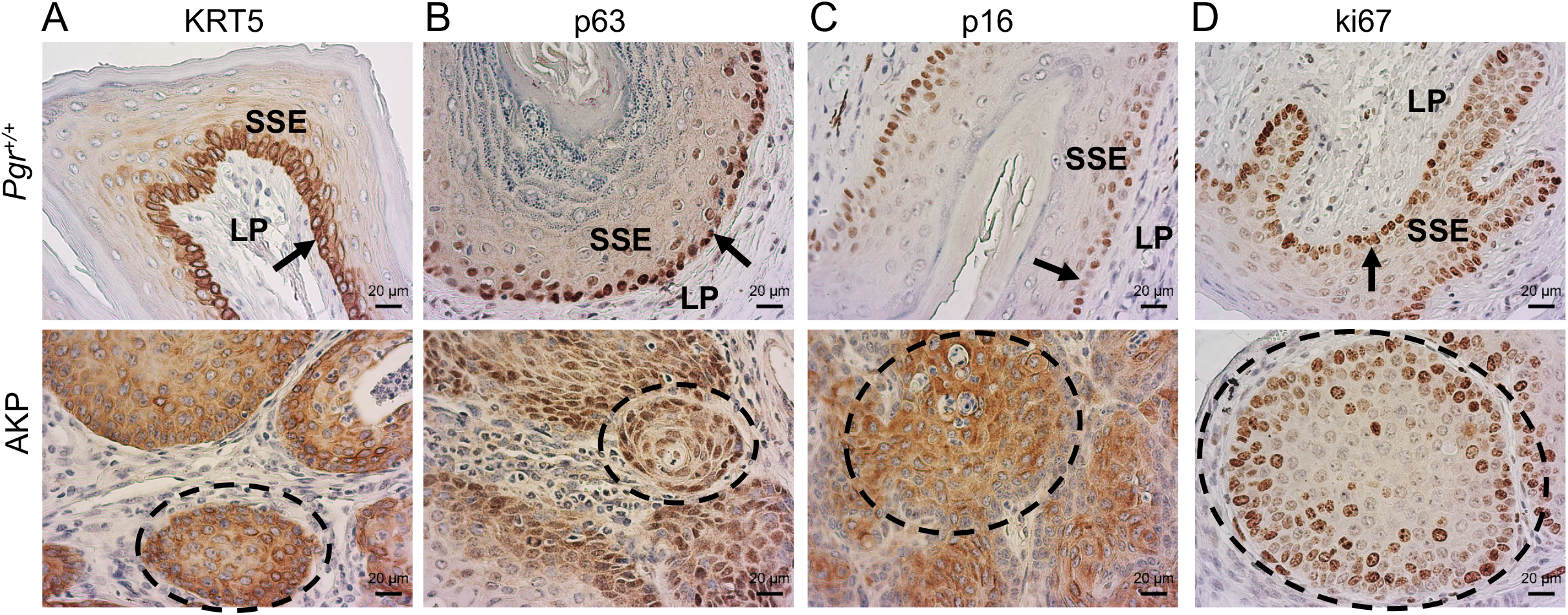
Molecular Immunohistochemical staining for (A) cytokeratin 5 (KRT5), (B) tumor protein P63 (p63), (C) cyclin dependent kinase inhibitor 2A (p16), and (D) marker of proliferation Ki-67 (ki67). SSE = stratified squamous epithelium, LP = lamina propria, arrow = basal layer, dashed circle = nests of squamous cell carcinoma. Scale bars, 20 μm.

### Hormone-dependent and estradiol responsive tumors

Ovaries were removed at six weeks (ov’ex), and AKP and *Pgr^+/+^* mice were examined six weeks later (age = 12 weeks). Ovarian hormone depleted AKP and *Pgr^+/+^* mice at 12 weeks had zero gross lesions (*n* = 8) (Supplementary Table S5), suggesting that AKP vaginal squamous cell carcinoma may be hormone dependent. To assess hormone responsiveness, ov’ex AKP and ov’ex *Pgr^+/+^* mice were treated with a 17β-estradiol pellet or sham for 60 days. Both ov’ex *Pgr^+/+^*-sham and ov’ex AKP-sham had no gross vaginal or uterine lesions, and the uteri were of grossly smaller size than intact mice (Figure 4; ov’ex *Pgr^+/+^*-sham: 0.51 +/-0.06 mg/g; *Pgr^+/+^* intact: 3.24+/-0.29 mg/g; two-tailed Student t-test; *P* <0.0001; and ov’ex AKP-sham: 0.48+/-0.054 mg/g; AKP intact: 4.25+/-0.29 mg/g; two-tailed Student t-test; *P* <0.0001). Ov’ex *Pgr^+/+^-E2* uteri were translucent and cystically dilated, consistent with the edematous effects of estradiol in the uterus, and ov’ex *Pgr^+/+^*-E2 mice exhibited no gross vaginal lesions (Figure 4). Ov’ex AKP-E2 uteri did not exhibit the characteristic cystic enlargement of estradiol treatment. Ov’ex AKP-E2 uteri were larger than ov’ex AKP-sham uteri (ov’ex AKP-sham: 0.48+/-0.054 mg/g; ov’ex AKP-E2: 24.6+/-15.2 mg/g; two-tailed Student t-test; *P* <0.0001). However, the ov’ex AKP-E2 uteri were more similar to AKP-intact uteri in that they appeared smooth and solid. The majority of ov’ex AKP-E2 mice (4 out 6) exhibited exophytic gross vaginal lesions (Figure 4 and Supplementary Table S6).

**Figure 4.**
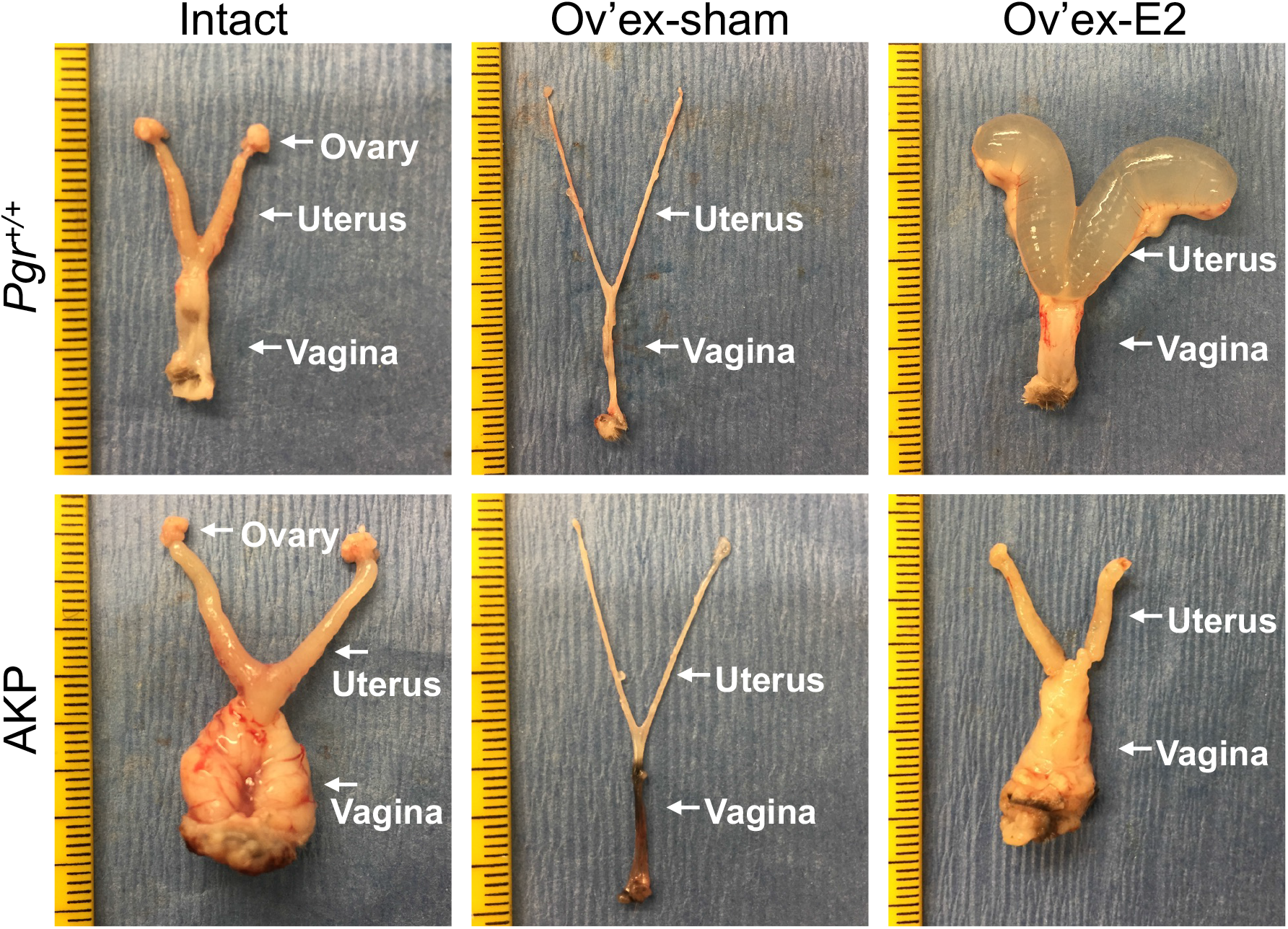
Gross lesions in AKP mice with 17β-estradiol treatment for 60 days. Intact AKP mice exhibited large gross vaginal lesions. Both ov’ex *Pgr^+/+^-sham* and ov’ex AKP-sham mice exhibited tiny, thin uteri without gross vaginal lesions. Ov’ex *Pgr^+/+^-E2* mice had enlarged fluid-filled uteri. Ov’ex AKP-E2 mice exhibited gross vaginal lesions but without the edematous cystic nature of the uterus. Ruler, tick mark indicates 1 mm.

Both ov’ex *Pgr^+/+^*-sham and ov’ex AKP-sham uteri were smaller than intact uteri (Supplementary Figure S9). Histologically, uteri from AKP intact mice showed endometrial hyperplasia with increased surface area of luminal epithelium, crowding of glandular epithelium, and little stroma between glands (Figure 5 and Supplementary Figure S9 and Supplementary Table S3). Neither ov’ex *Pgr^+/+^*-sham nor ov’ex AKP-sham showed any evidence of endometrial hyperplasia. Both contained a single layer of benign luminal epithelium and normal number of endometrial glands without evidence of atypia (Figure 5 and Supplementary Figure S9 and Supplementary Table S6).

**Figure 5.**
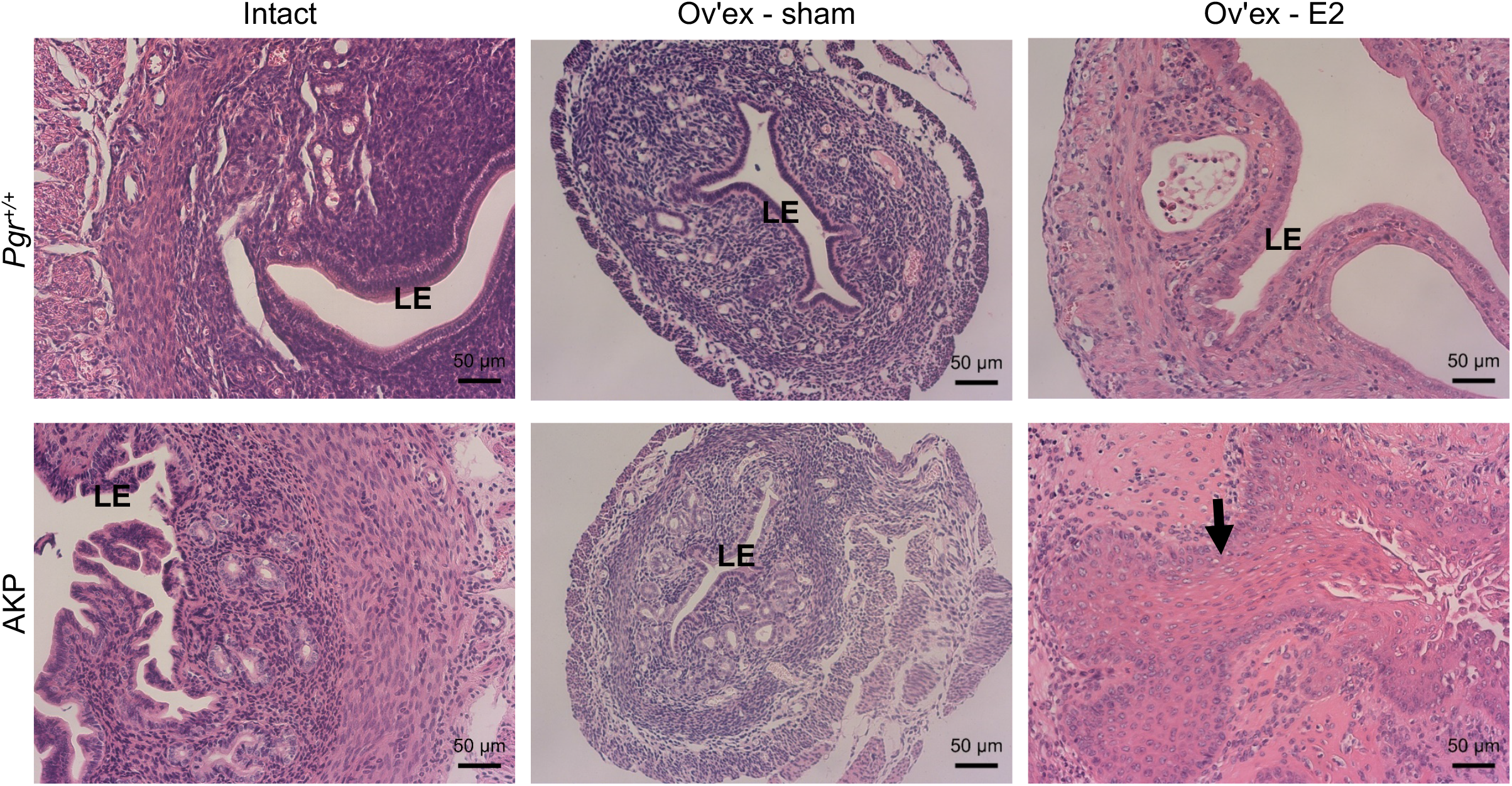
Uterine histology in *Pgr^+/+^* and AKP mice with and without 17β-estradiol (E2) treatment. 16-week-old intact *Pgr^+/+^* and AKP mouse uteri are shown for comparison. AKP intact uteri shown exhibits endometrial hyperplasia. Ov’ex AKP-sham are similar in histology to ov’ex *Pgr^+/+^-sham.* Ov’ex *Pgr^+/+^-E2* exhibited enlarge dilated uteri without evidence of endometrial hyperplasia or atypia. Ov’ex AKP-E2 exhibited squamous metaplasia of the endometrium (black arrow). LE = luminal epithelium. Scale bars, 50 μm. Hematoxylin and eosin staining.

Ov’ex *Pgr^+/+^* mice treated with 17β-estradiol for 60 days (ov’ex *Pgr^+/+^-E2)* showed edematous, enlarged uteri. Clear fluid resulted in dilated lumina. The luminal epithelium existed in a single layer that maintained polarity without evidence of nuclear atypia (Figure 5 and Supplementary Figure S9). Ov’ex AKP-E2 uteri were not dilated by fluid. Instead, they were largely lined by epithelium with squamous metaplasia without evidence of adenocarcinoma or nuclear atypia (Figure 5 and Supplementary Figure S9). Other non-malignant phenotypes included endometrial hyperplasia (1/6 mice) and stump pyometra (2/6 mice) (Supplementary Table S6). The hyperplasia and metaplasia seen in ov’ex AKP-E2 mice (4/6) was more penetrant than the hyperplasia observed in intact 16-week mice (3/11).

The vaginal tissue was also hormone responsive. Histologically, hormone depletion in *Pgr^+/+^* vaginas (ov’ex *Pgr^+/+^*-sham) resulted in a thin single epithelial layer, with decreased keratinization (Figure 6 and Supplementary Figure S10), consistent with models of vaginal atrophy in rodents [40]. Treatment with 17β-estradiol pellets for 60 days (ov’ex *Pgr^+/+^-E2)* reversed these effects similar to previous studies [40]. Ov’ex *Pgr^+/+^*-E2 vaginas exhibited normal keratinized stratified squamous epithelium and normal lamina propria (Figure 6 and Supplementary Figure S10). Similarly, ov’ex-AKP-sham vaginas exhibited a thin layer of vaginal epithelium although it appeared more than a single layer. Treatment with estradiol reversed the benign, atrophic vaginal histology in the ov’ex AKP vaginas to squamous cell carcinoma (Figure 6 and Supplementary Figure S10). Histological description of each animal can be found in Supplementary Table S6.

**Figure 6.**
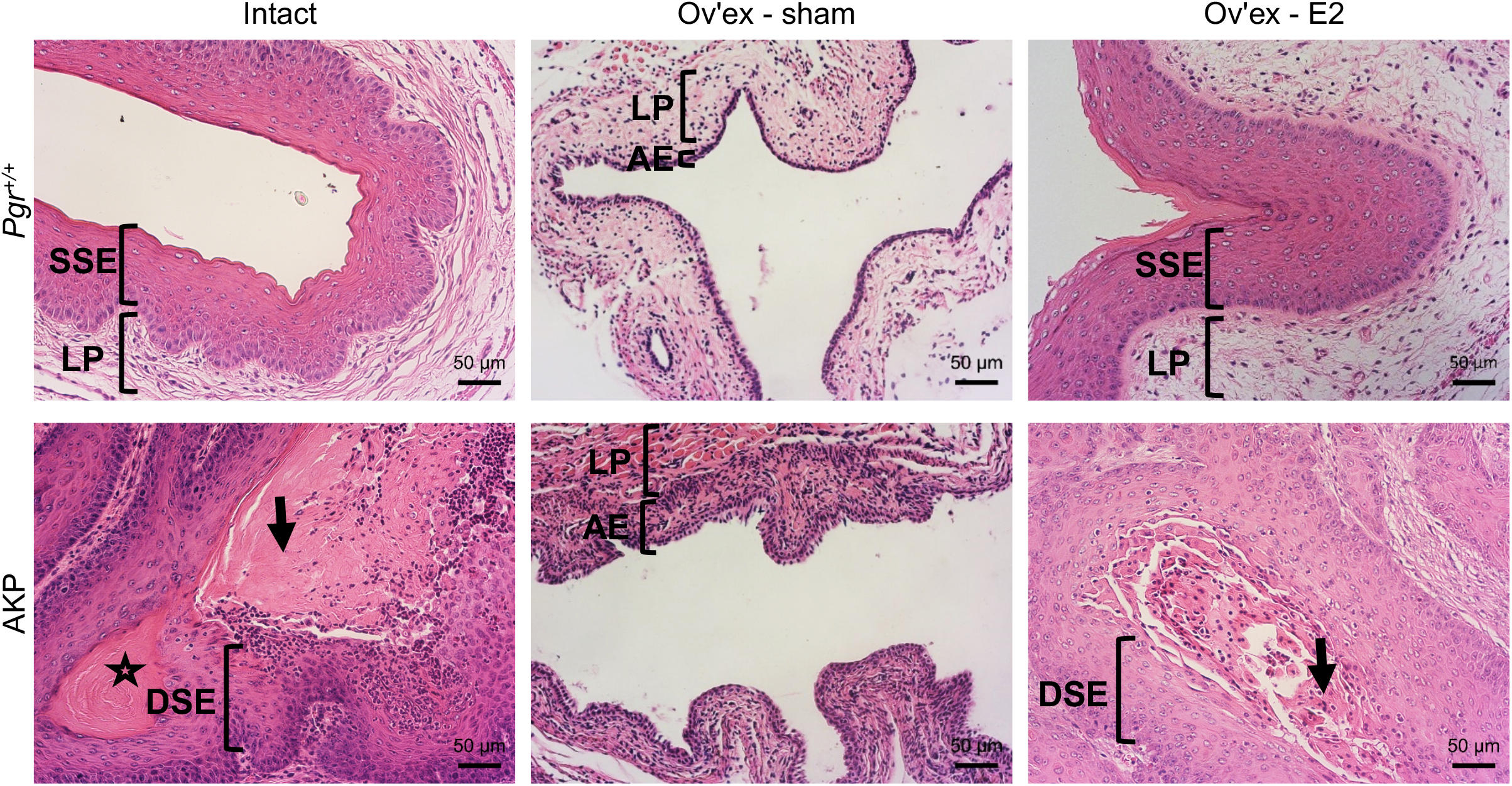
Vaginal histology in *Pgr^+/+^* and AKP mice with and without 17β-estradiol (E2) treatment.16-week-old intact *Pgr^+/+^* and AKP mouse vaginas are shown for comparison. Intact *Pgr^+/+^* vaginas exhibited normal keratinized stratified squamous epithelium (SSE) with underlying lamina propria (LP). Intact AKP vaginas show squamous cell carcinoma with central keratinization (star), eosinophilic cytoplasm (arrow), and dysplastic squamous epithelium (DSE). Ov’ex *Pgr^+/+^*-sham vaginas exhibited signs of vaginal atrophy with thin atrophic epithelium (AE) and thin lamina propria. Hormone replacement resulted in a thickened epithelial layer with keratinization and normal lamina propria. Ov’ex AKP-sham mice exhibited thinning of vaginal epithelium without evidence of carcinoma. Ov’ex AKP-E2 mice exhibited squamous cell carcinoma with dysplastic squamous epithelium (DSE) and eosinophilic cytoplasm (black arrow). Scale bars, 50 μm. Hematoxylin and eosin staining.

## DISCUSSION

A small subset of mutated genes are enriched in female reproductive tract malignancies [4], but the functional role of each gene is unclear. To study two such genes, we created the AKP mouse. In our study, AKP mice develop highly penetrant, hormone-responsive squamous cell carcinoma in the vagina. Of note the vaginal squamous cell carcinoma described here is human papillomavirus-(HPV) independent. Importantly, TCGA studies showed that HPV-independent cervical cancer may have ~6% *KRAS* mutations and 30% have loss-of-function mutations in ARID1A [11]. Therefore, this model potentially represents a unique molecular subset of vaginal squamous cell carcinoma.

Because the *Pgr^Cre^* mouse is a powerful Cre recombinase for endometrial cancer modeling [41], our original hypothesis was that AKP female mice would develop endometrial cancer. Oncogenic *KRAS* has been detected in up to 30% of endometrial cancers [18–20]. More than 40% of endometrial cancers have a mutation in *ARID1A* [7–9]. Loss of a proposed tumor suppressor and gain of an oncogene in the uterus should have led to endometrial cancer. The aggressive nature of the vaginal tumors leading to early euthanasia for humane reasons may explain the malignancy limited to the vagina rather than the endometrium. By 16 weeks AKP mice had highly aggressive vaginal tumors limiting the ability to study other gynecological malignancies that may develop at older time points. To support this notion, the hyperplasia and epithelial cell squamous metaplasia seen in ov’ex AKP-E2 mice (4/6) was more penetrant than the hyperplasia observed in intact 16-week AKP females (3/11). However, even with long-term (60 day) 17β-estradiol treatment, endometrial cancer was not discovered, suggesting that something molecular may be restraining cancer in the endometrium rather than the vagina. Similarly, in the pancreatic cancer field, oncogenic *Kras* was found to be insufficient to drive pancreatic adenocarcinoma due to the protective effects of ARID1A [42–47]. For example, oncogenic expression of *Kras^G12D^* alone leads to pancreatic intraepithelial lesions but not malignancy. Following a loss of function mutation in *Arid1a,* the *Kras^G12D^-*expressing mice developed pancreatic ductal adenocarcinoma. These studies all support the hypothesis that ARID1A restrains malignant transformation from benign lesions driven by oncogenic *Kras^G12D^.* Work in pancreatic adenocarcinoma [42–47] has identified escape mechanisms that need to be explored in the endometrium.

Mechanistically, the early development of aggressive and highly penetrant vaginal tumors may be in part due to the suppression of oncogenic-induced cellular senescence mediated by ARID1A loss. Cellular senescence plays a vital role in vaginal development; in fact, when senescence is blocked during murine development mice generate septate vaginas [48]. Cellular senescence is frequently found in aging or cancerous tissue, potentially due to oncogenic signaling [49]. Oncogenic KRAS^G12D^ expression in pancreatic cancer cell lines induces cellular senescence [47]. Of note, ARID1A knockdown repressed oncogenic-induced cellular senescence in KRAS^G12D^ pancreatic cell lines, leading to cell cycle progression [47]. Thus, we posit that deletion of *Arid1a* with gain of oncogenic *Kras^G12D^* leads to loss of oncogene-induced cellular senescence and unchecked proliferation and malignant transformation. Consistent with this view, ARID1A promoter hypermethylation and decreased ARID1A expression led to increase progression of squamous cell carcinoma *in vitro* and *in vivo* [50]. CDKN2A or p16 is potential marker of cellular senescence. Specifically, p16 staining was found, as expected [36,39], in the *Pgr^+/+^* mouse localized to the nucleus. The AKP vaginal squamous cell carcinoma expressed p16 in both the nucleus and the cytoplasm (Figure 3). Cytoplasmic p16 staining was previously considered an artifact, although there is evidence that cytoplasmic staining is tumor specific [51,52]. In fact, it was hypothesized that cytoplasmic staining represents the cellular mechanism in which p16 becomes inactivated and allows for tumor progression [51,52]. Cytoplasmic p16 has been studied as a potential prognostic marker in squamous cell carcinoma of the head and neck, with increased survival correlating with higher cytoplasmic staining [53–55]. Both the functional role of oncogenic cellular senescence in vaginal squamous cell carcinoma development and the mechanistic role of *Arid1a* in the translocation of p16 need further investigation.

In humans, *Kras* mutations were not found in vaginal squamous cell carcinoma [4], but *Kras^G12D-^* induced vaginal lesions have been previously reported in mice [24,56]. Kim *et al.* and Gades *et al.* both reported vaginal papilloma when *Kras^G12D^* was mutated through a progesterone or insulin promoter factor mediated-Cre. These lesions had high penetrance but were non-malignant [24,56]. Their benign nature suggests that a secondary hit may be necessary for transformation to malignancy. This hypothesis is supported by the presence of vaginal squamous cell carcinoma in *Kras^G12D^* mice with a *Pten* inactivating mutation driven by vaginal delivered adenovirus-Cre [25]. Blum *et al.* described the vaginal lesions as exophytic protruding externally from the vagina and histologically resembled those described here in AKP mice [25]. Similarly, AKP female mice exhibited vaginal tumors that were malignant earlier (Figure 2) than KP female mice developed non-malignant lesions [24].

The effects of 17β-estradiol treatment on tumor growth were surprising. Clinically, vaginal squamous cell carcinoma is not treated as a hormone-responsive disease. While clinically 17β-estradiol is used to increase epithelial cell thickness and treat the symptoms associated with postmenopausal vaginal atrophy, the mechanisms involved in squamous cell proliferation and differentiation are not well studied. Further, the effects of 17β-estradiol treatment on progesterone receptor expression and thus, function of Cre recombinase in the vagina need further study.

Clinically, 17,600 women were diagnosed with primary vaginal squamous cell carcinoma and 8062 women died from vaginal squamous cell carcinoma worldwide in 2018 [3]. Therefore, the continued development of new models is critical to understanding the mechanistic changes. As the impact of the HPV vaccine decreases the prevalence of HPV-dependent squamous cell carcinoma, this model of HPV-independent squamous cell carcinoma will be even more important. We have described a genetically engineered mouse model that targets the knockout of the tumor suppressor *Arid1a* and the knock in of oncogenic *Kras^G12D^* in the gynecologic tract, leading to the development of primary squamous cell carcinoma of the vagina. We describe the progression of disease both within the vagina and benign lesions found outside the vagina. We highlighted the role of hormone expression in primary squamous cell carcinoma of the vagina, alluding to potential mechanistic and therapeutic targets. Finally, we proposed a potential mechanism for tumor progression of repressed oncogenic-induced cellular senescence via translocation of p16 from the nucleus to the cytoplasm. This genetically engineered mouse model of primary squamous cell carcinoma of the vagina may lead to advances in early detection, understanding of initiation and progression, and novel treatment options.

## Acknowledgements

We thank the Indiana Center for Musculoskeletal Health Histology Core at Indiana University School of Medicine and the Human Tissue and Acquisition and Pathology Core at the Dan L. Duncan Comprehensive Cancer Center at Baylor College of Medicine for histology services, and University of Virginia Center for Research in Reproduction Ligand Assay and Analysis Core for the hormone assay.

The University of Virginia Center for Research in Reproduction Ligand Assay and Analysis Core is supported by the Eunice Kennedy Shriver NICHD Grant R24 HD102061. This work was supported by The Liz Tilberis Scholarship Ovarian Cancer Research Fund through the Estate of Agatha Fort (to S.M.H.), 1R03 CA19127 A1 (to SMH), NIH/NICHD RO1 HD042311 (to JPL), and Coordenação de Aperfeiçoamento de Pessoal de Nível Superior -(CAPES) - Brazil (to MSLP). The Intramural Research Program of the National Institute of Environmental Health Sciences supported FJD: Project Z1AES103311-01. We thank Julio Agno for technical support. We thank Dr. Joanne S. Richards for kindly providing the *Kras^Lox-Stop-Lox-G12D/+^* mice and Dr. Zhong Wang for kindly providing the *Arid1a^flox/flox^* mice.

## Statement of Author Contributions

XW, MP, and SMH contributed to study design, data collection, data analysis, data interpretation, literature search, and generation of figures. JRHW contributed to data interpretation, literature search, and generation of figures. REE contributed to data analysis, data interpretation, literature search, and generation of figures. JPL and FJD contributed new reagents/analytic tools. All authors were involved in writing and editing the paper and had final approval of the submitted and published versions.

Supplementary Figures document as PDF contains figure legends with supplementary figures.

Supplementary Tables S1-S6 are included in the Excel sheet.

