## Supplementary Figures S1-S10 for "Vaginal squamous cell carcinoma develops in mice with *Arid1a* loss and gain of oncogenic *Kras*"

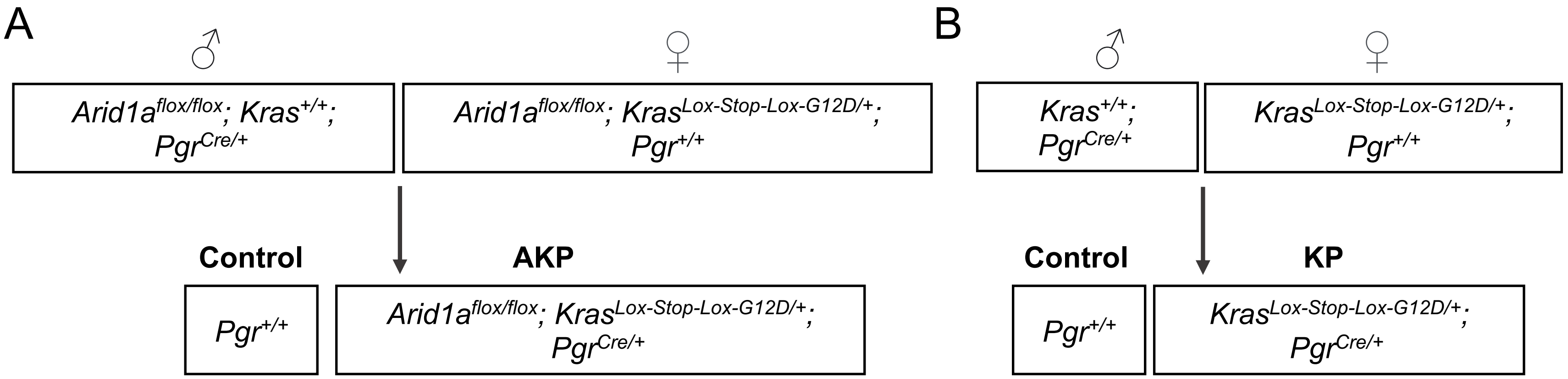


C

| Primer Name | Primer Sequence |
|---|---|
| *Pgr Cre* P1 | 5'- TATACCGATCTCCCTGGACG -3' |
| *Pgr Cre* P2 | 5'- ATGTTTAGCTGGCCCAAATG -3' |
| *Pgr Cre* P3 | 5'- CCCAAAGAGACACCAGGAAG -3' |
| *Kras* F | 5'- AGCCACCATGGCTTGAGTAA -3' |
| *Kras* R | 5'- CTTTACAAGCGCACGCAGAC -3' |
| *Arid1a* F | 5'- GTAATGGGAAAGCGACTACTGGAG -3' |
| *Arid1a* R | 5'- TGTTCATTTTTGTGGCGGGAG -3' |

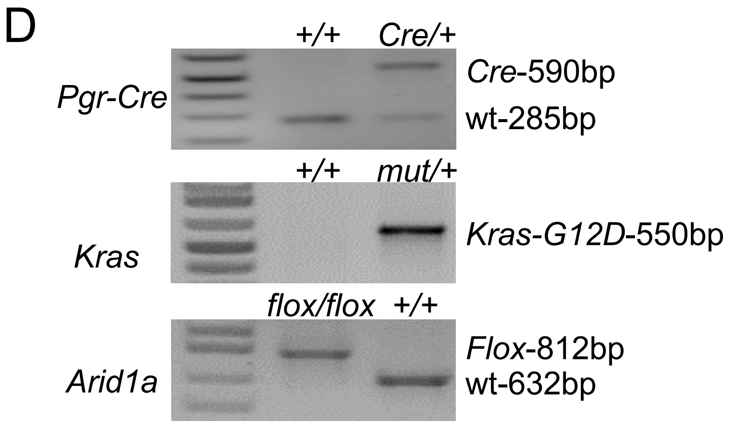


**Supplementary Figure S1.** Generation of experimental mice. Final breeding strategy for generation of (A) AKP (*Arid1a*$^{flox/flox}$*; Kras*$^{Lox-Stop-Lox-G12D/+}$*; Pgr*$^{Cre/+}$), (B) KP (*Kras*$^{Lox-Stop-Lox-G12D/+}$*; Pgr*$^{Cre/+}$), and control female mice. The *Cre* allele was maintained in male mice for breeding. (C) PCR primers for genotyping. (D) Tail genotyping showing each *Cre*, floxed, mutated (mut), or wild-type (wt) allele.

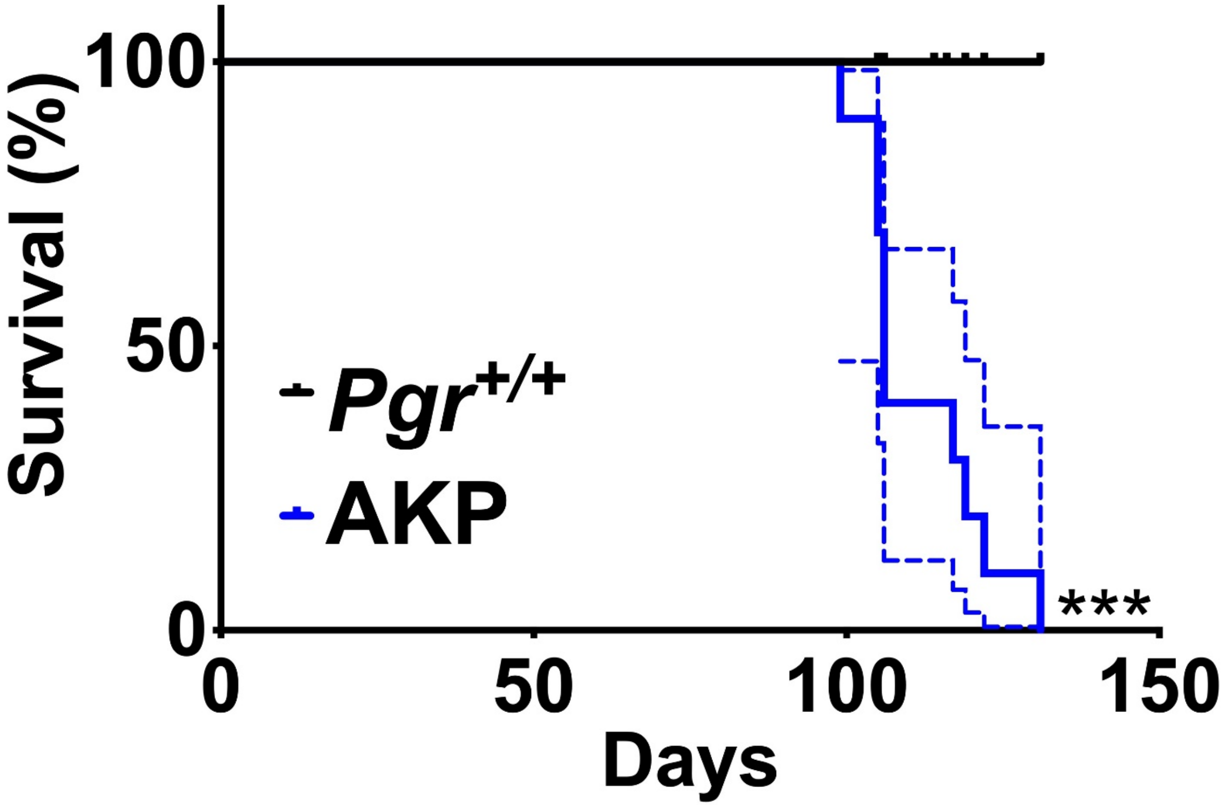


**Supplementary Figure S2**. AKP mice have decreased survival compared to control mice. Kaplan-Meier survival curves were analyzed by log-rank (Mantel-Cox) test. Dashed lines indication 95% confidence intervals. *** $P < 0.001$. $n = 10$ per group.

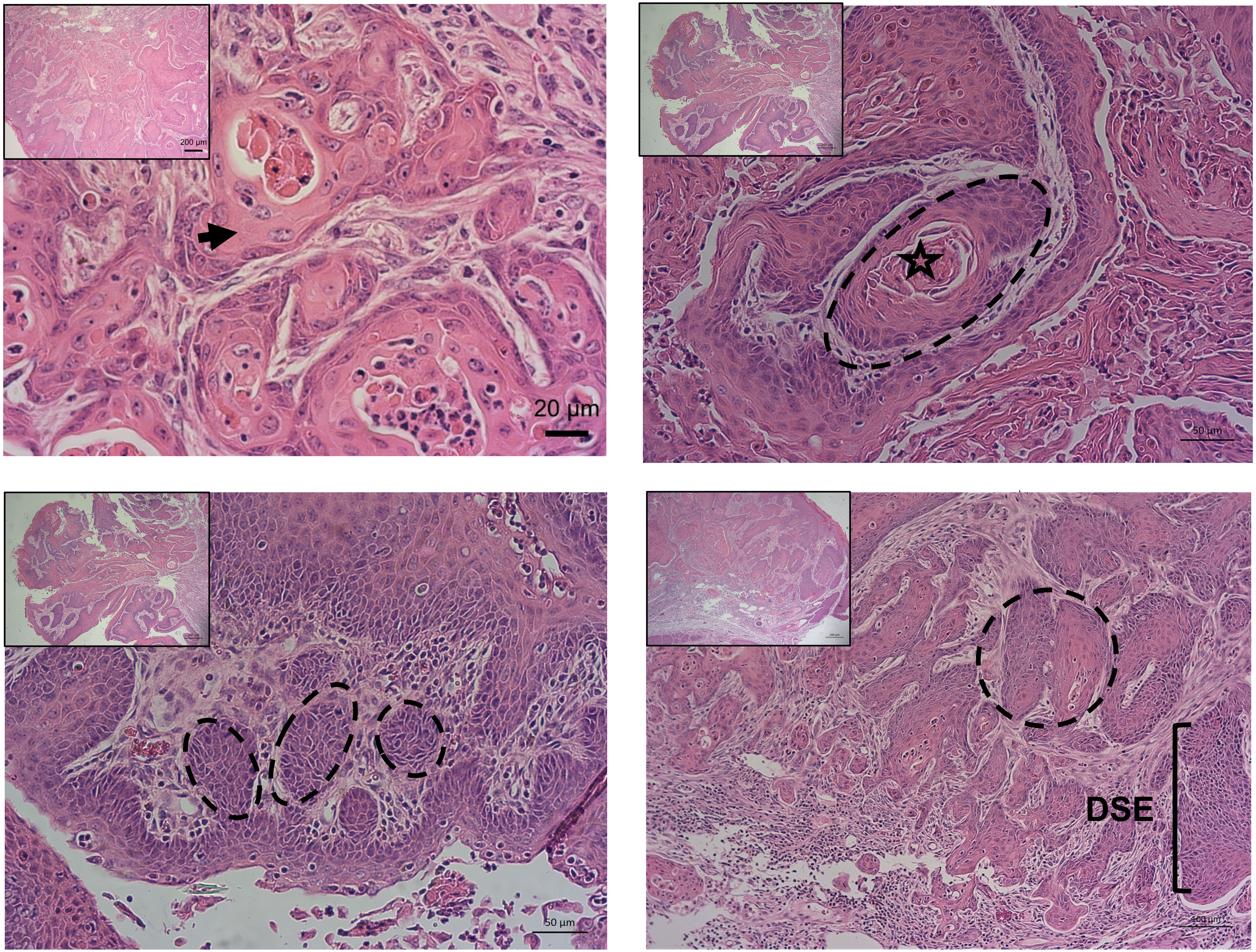


**Supplementary Figure S3**. Additional images of invasive squamous cell carcinoma in AKP mice with nest of cells (dashed circles), abundant eosinophilic cytoplasm (black arrow), central keratinization (star), and dysplastic squamous epithelium (DSE). Scale bars, low-power, 200 µm; high-power, 20-100 µm. Hematoxylin and eosin staining.

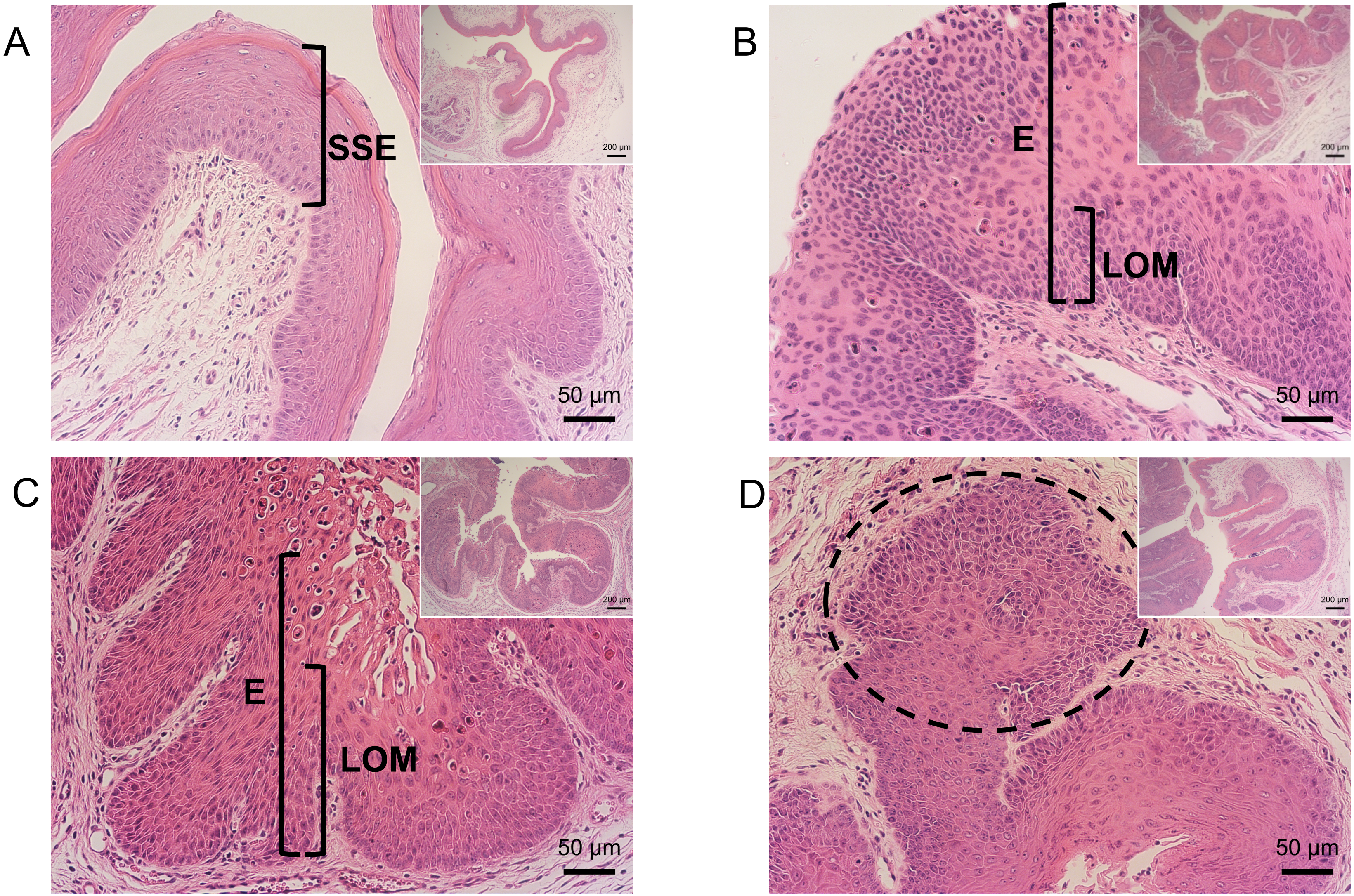


**Supplementary Figure S4**. Squamous intraepithelial lesions and squamous cell carcinoma in AKP mice at 8 weeks. (A) *Pgr*$^{+/+}$ vagina showed normal stratified squamous epithelium (SSE) with normal keratinization. AKP vaginas contained (B) LSIL with increased nuclear-to-cytoplasmic ratios and loss of maturation (LOM) confined to the lower third of the epithelium (E), (C) HSIL with cell crowding, high nuclear-to cytoplasmic ratio, and loss of maturation in at least two-thirds of the epithelium, and (D) squamous cell carcinoma. Scale bars, low-power (inset), 200 µm; high-power, 50 µm. Hematoxylin and eosin staining.

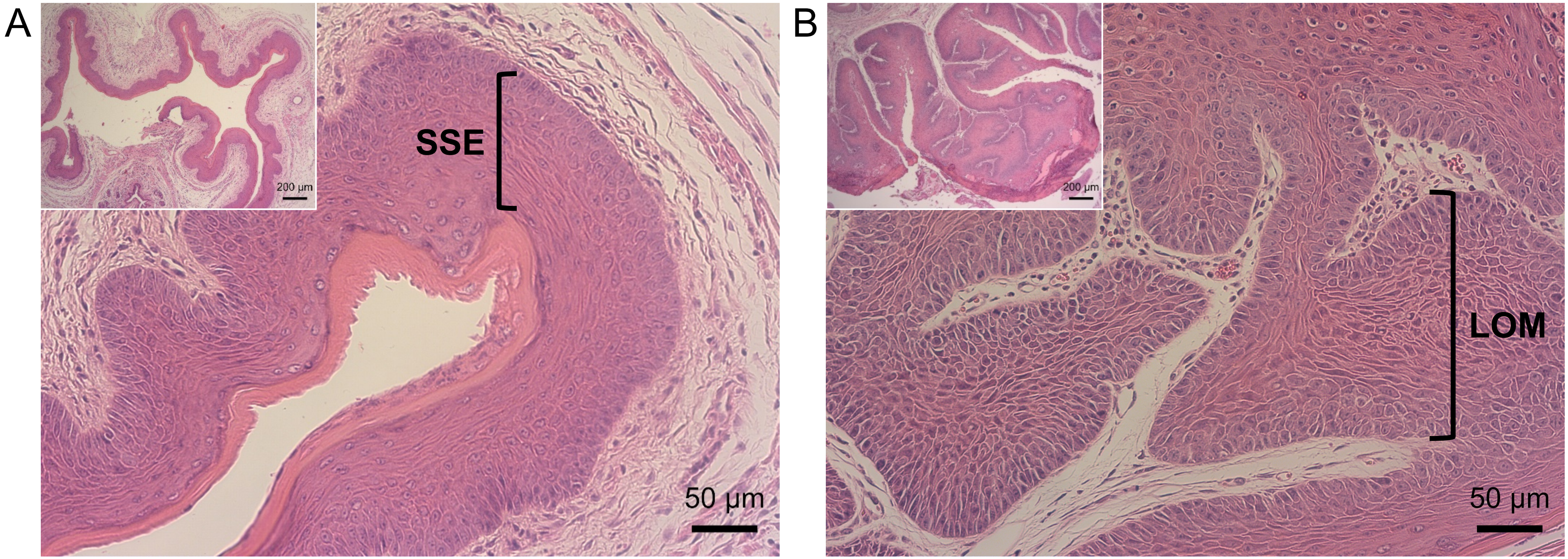


**Supplementary Figure S5**. Non-malignant vaginal lesions in 8-week KP animals. (A) Normal vagina of a *Pgr*$^{+/+}$ mouse at 8 weeks with stratified squamous epithelium (SSE) and normal keratinization. (B) HSIL from KP mouse with high nuclear-to cytoplasmic ratio and loss of maturation (LOM) in at least two-thirds of the epithelium. Scale bars, low-power (inset), 200 µm; high-power, 50 µm. Hematoxylin and eosin staining.

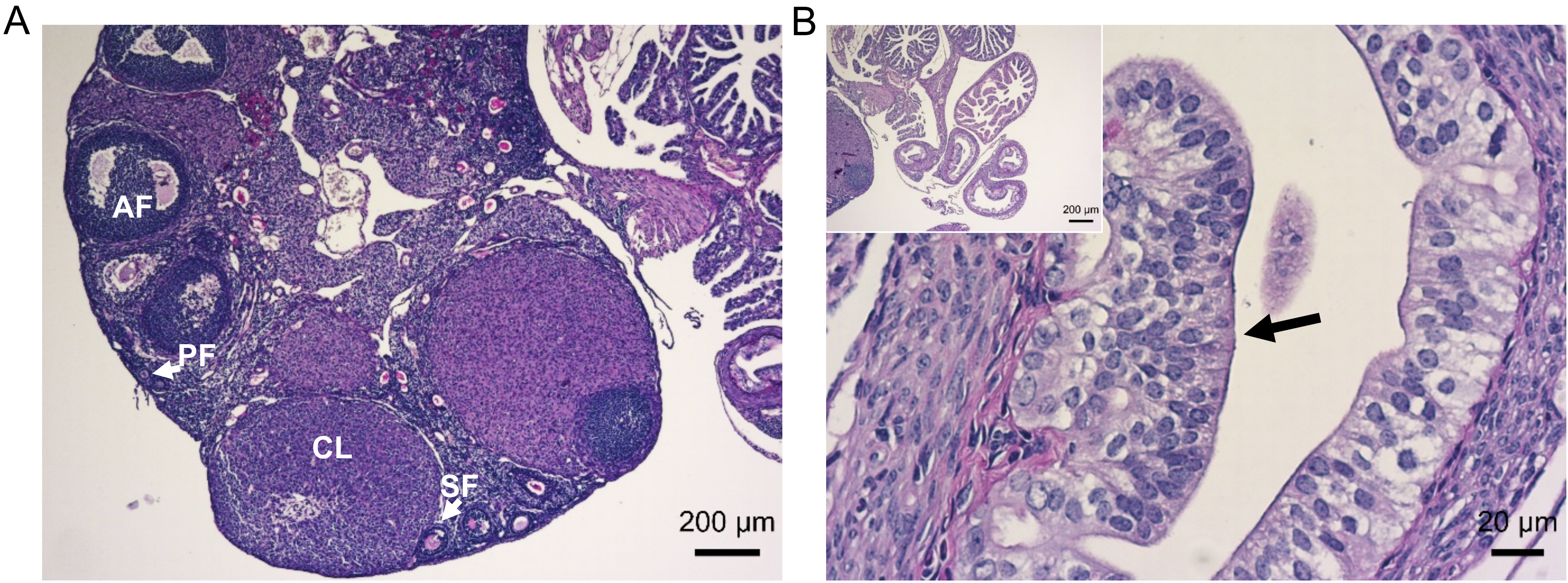


**Supplementary Figure S6**. AKP adnexal histology. (A) The AKP mice ovaries showed normal histology with follicles in each stage of follicular development. There were primary (PF), secondary (SF), and antral (AF) follicles and corpus luteum (CL). Scale bar, 200 µm. (B) Some AKP mice oviducts showed atypical epithelium as evidenced by >1 cell thickness epithelium (black arrow). Scale bars, low-power (inset), 200 µm; high-power, 20 µm. Periodic acid-Schiff staining.

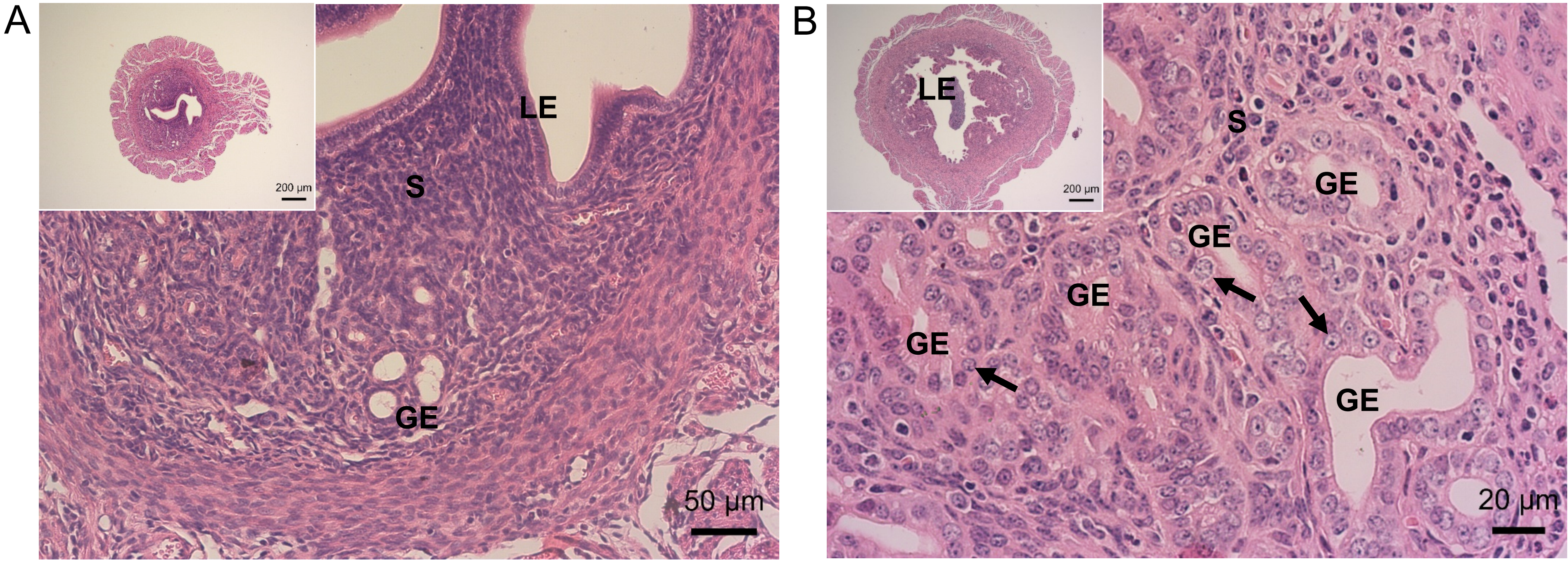


**Supplementary Figure S7**. Endometrial phenotype in AKP uterus. (A) Pathologically normal *Pgr*$^{+/+}$ mouse uterus with normal glandular epithelium (GE), luminal epithelium (LE), and adequate intervening stroma (S). (B) Endometrial hyperplasia in AKP mice with nuclear atypia in the endometrial glandular epithelium (GE) at 16-weeks. Nuclear atypia was identified by rounding of nuclei (arrow) and loss of polarity. Scale bars, low-power (inset), 200 µm; high-power, (A) 50 µm and (B) 20 µm. Note the difference in size between control inset (A) and AKP inset (B) at same magnification. Hematoxylin and eosin staining.

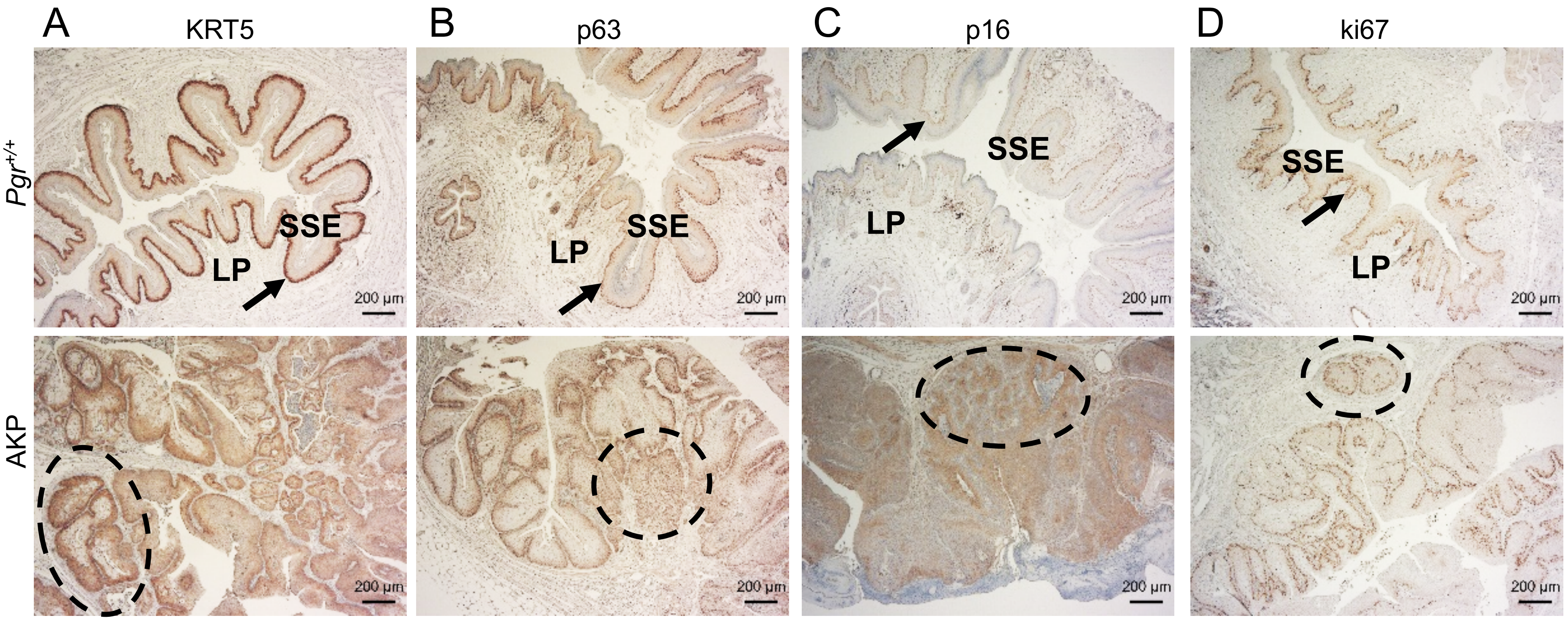


**Supplementary Figure S8**. Immunohistochemical staining for (A) cytokeratin 5 (KRT5), (B) tumor protein P63 (p63), (C) cyclin dependent kinase inhibitor 2A (p16), and (D) marker of proliferation Ki-67 (ki67). SSE = stratified squamous epithelium, LP = lamina propria, arrow = basal layer, dashed circle = nests of squamous cell carcinoma. These images represent lower magnification views of Figure 3. Scale bars, 200 µm.

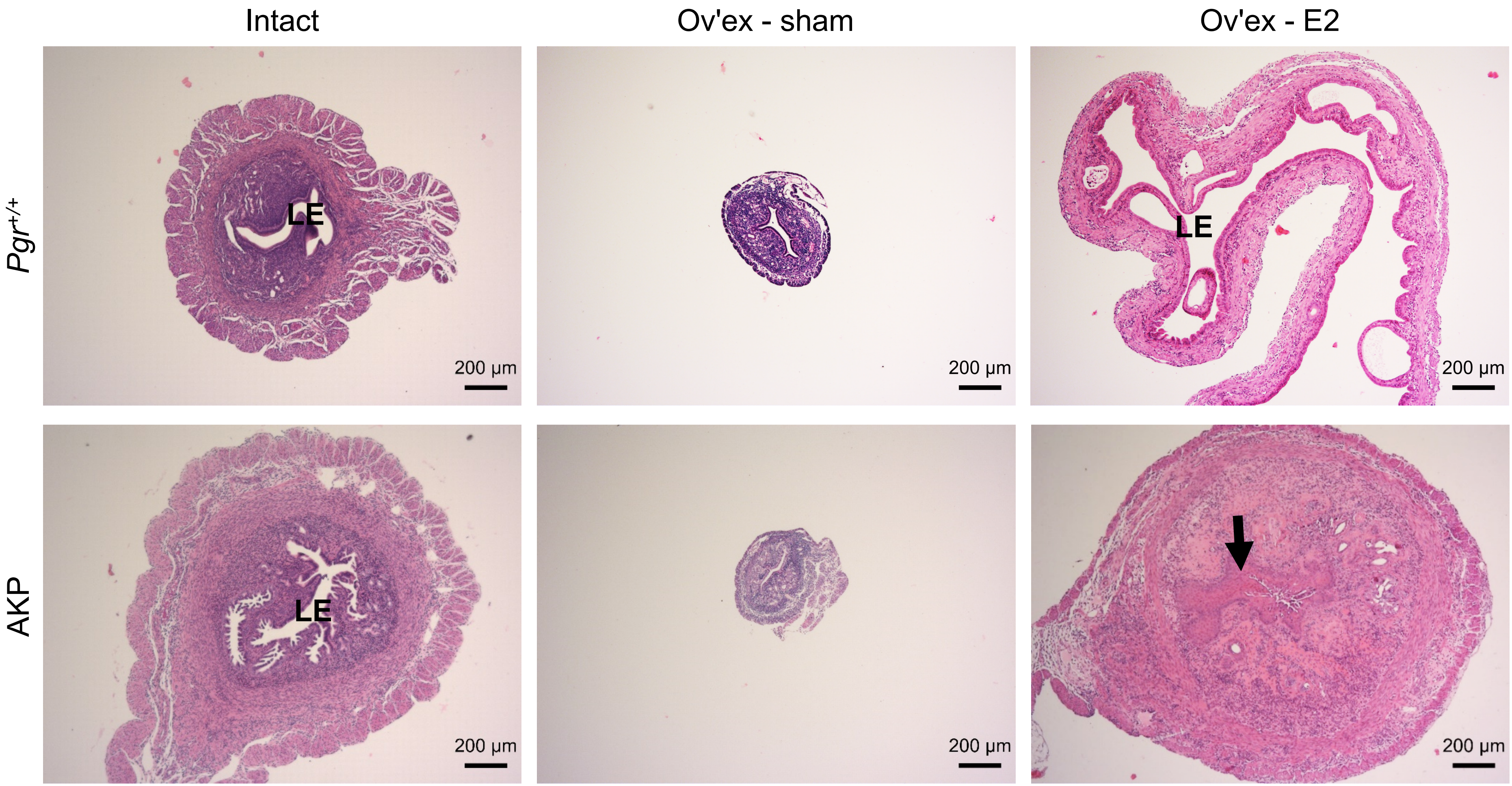


**Supplementary Figure S9**. Uterine histology in *Pgr*$^{+/+}$ and AKP mice with and without 17β-estradiol treatment. 16-week-old intact *Pgr*$^{+/+}$ and AKP mouse uteri are shown for comparison. AKP intact uteri exhibit endometrial hyperplasia. Ov'ex AKP-sham are similar in histology to ov'ex *Pgr*$^{+/+}$-sham. Ov'ex *Pgr*$^{+/+}$-E2 exhibited enlarge dilated uteri without evidence of endometrial hyperplasia or atypia. Ov'ex AKP-E2 exhibited squamous metaplasia (black arrow). LE = luminal epithelium. These images represent lower magnification views of Figure 5. Scale bars, 200 µm. Hematoxylin and eosin staining.

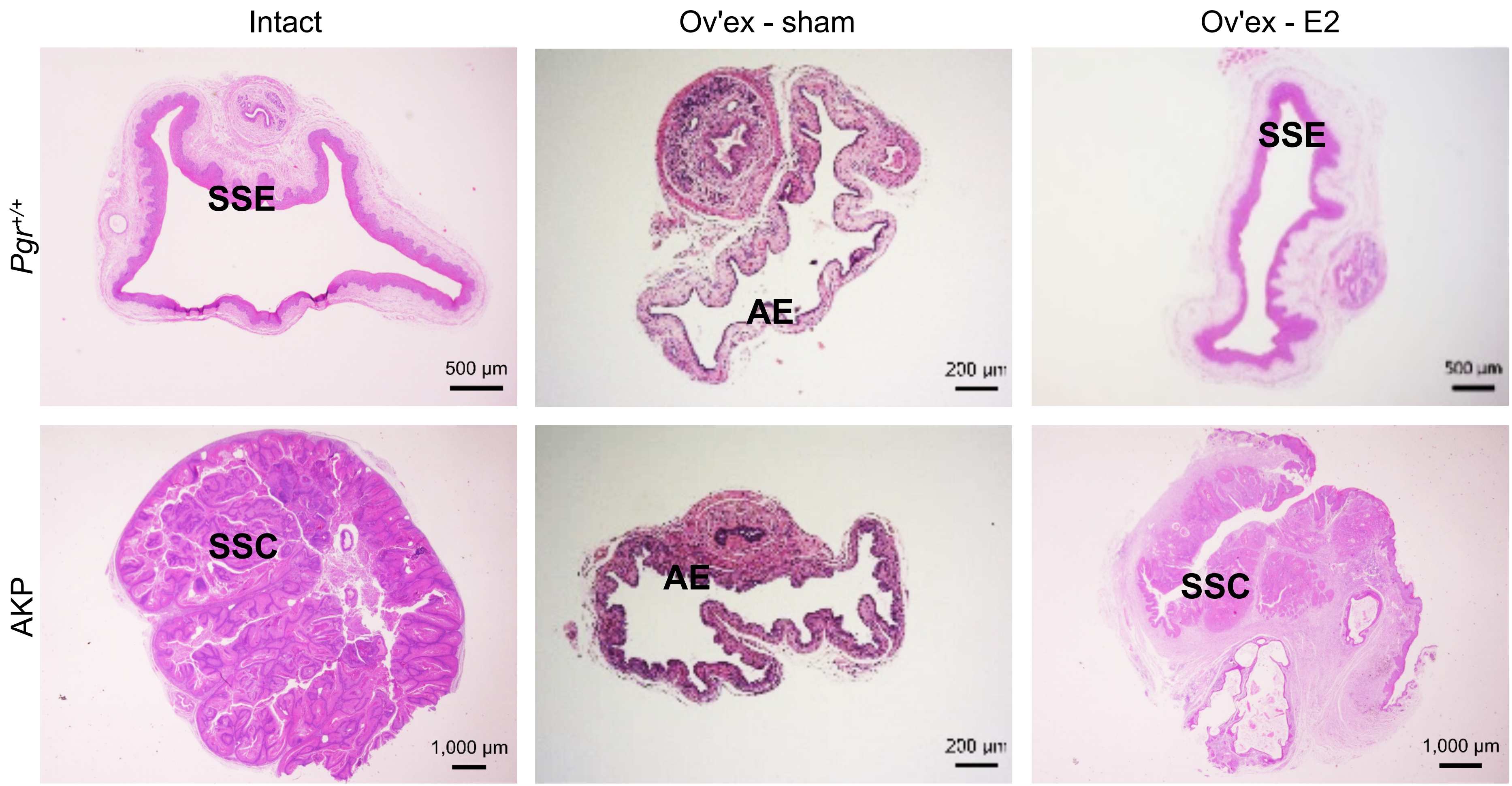


**Supplementary Figure S10**. Vaginal histology in *Pgr*$^{+/+}$ and AKP mice with and without 17β-estradiol (E2) treatment. 16-week-old intact *Pgr*$^{+/+}$ and AKP mouse vaginas are shown for comparison. Intact *Pgr*$^{+/+}$ vaginas exhibited normal keratinized squamous cell epithelium (SSE) and lamina propria. Intact AKP vaginas show squamous cell carcinoma (SSC) with carcinoma encompassing the entire vaginal opening. Ov'ex *Pgr*$^{+/+}$-sham vaginas exhibited signs of vaginal atrophy with thin atrophic epithelium (AE) and thin lamina propria. Hormone replacement resulted in a thickened epithelial layer with keratinization and normal lamina propria. Ov'ex AKP-sham mice exhibited thinning of vaginal epithelium without evidence of carcinoma. Ov'ex AKP-E2 mice exhibited squamous cell carcinoma. Scale bars, 200 µm, 500 µm, or 1000 µm. Hematoxylin and eosin staining.
